# Revealing the Grammar of Small RNA Secretion Using Interpretable Machine Learning

**DOI:** 10.1101/2023.04.04.535452

**Authors:** Bahar Zirak, Mohsen Naghipourfar, Ali Saberi, Delaram Pouyabahar, Amirhossein Zarezadeh, Lixi Luo, Lisa Fish, Doowon Huh, Albertas Navickas, Ali Sharifi-Zarchi, Hani Goodarzi

## Abstract

Small non-coding RNAs can be secreted through a variety of mechanisms, including exosomal sorting, in small extracellular vesicles, and within lipoprotein complexes ^1,2^. However, the mechanisms that govern their sorting and secretion are still not well understood. In this study, we present ExoGRU, a machine learning model that predicts small RNA secretion probabilities from primary RNA sequence. We experimentally validated the performance of this model through ExoGRU-guided mutagenesis and synthetic RNA sequence analysis, and confirmed that primary RNA sequence is a major determinant in small RNA secretion. Additionally, we used ExoGRU to reveal *cis* and *trans* factors that underlie small RNA secretion, including known and novel RNA-binding proteins, e.g., YBX1, HNRNPA2B1, and RBM24. We also developed a novel technique called exoCLIP, which reveals the RNA interactome of RBPs within the cell-free space. We used exoCLIP to reveal the RNA interactome of HNRNPA2B1 and RBM24 in extracellular vesicles. Together, our results demonstrate the power of machine learning in revealing novel biological mechanisms. In addition to providing deeper insight into complex processes such as small RNA secretion, this knowledge can be leveraged in therapeutic and synthetic biology applications.

## Introduction

Small non-coding RNAs play a variety of regulatory functions in the cell, including regulation of mRNA stability and protein synthesis ^3,4^. However, some small RNAs also reside in the extracellular space, packaged within extracellular vesicles or lipoprotein complexes, for example, where they are thought to play roles in cellular communication ^1,5,6^. Many recent studies have focused on the role of these secreted small RNAs as potential biomarkers in various diseases, particularly cancer^7–9^. RNA secretion, however, is not a random process. While some studies have focused on identifying the various mechanisms through which small RNAs are secreted ^10,11^, our knowledge of the underlying regulatory programs that govern extracellular sorting remains incomplete.

To reveal the *cis*-regulatory grammar that underlies small RNA secretion, we developed ExoGRU, a deep-learning model for predicting secretion probabilities of small RNAs based on their primary sequence. In addition to the commonly used machine learning performance metrics, we also used two independent experimental approaches to validate the veracity of our model. We used ExoGRU to (i) identify mutations that abrogate the secretion of known cell-free small RNAs, and (ii) predict high confidence sets of synthetic sequences that are secreted or retained. Having confirmed the accuracy of ExoGRU using these experimental strategies, we interrogated the model to reveal the cis-regulation RNA secretion grammar that it has learned. In addition to recapitulating known RNA binding proteins involved in small RNA sorting, such as YBX1, we also discovered and validated RBM24 as a novel RNA secretory factor. We also developed exoCLIP, a variation of CLIP-seq ^12^, that reveals RBP-RNA interactions in the cell-free space using UV cross-linking immunoprecipitation followed by high-throughput sequencing. Application of exoCLIP to RBM24 and HNRNPA2B1, another factor that was nominated by our model and previously implicated in RNA secretion, further confirmed their direct interactions with target small RNAs in extracellular vesicles.

Our results collectively show the significance of machine learning in uncovering previously unknown biological mechanisms. In addition to capturing the sequence features that mark small RNAs for secretion, our approach provides readily testable hypotheses around the key *trans* factors involved. This not only deepens our understanding of an intricate biological process, but also has practical implications for the design of artificial cell-free RNA species in synthetic biology applications.

## Results

### ExoGRU, A computational model for accurate prediction of small RNA secretion

To learn the small RNA secretory grammar, we first aggregated, curated, and labeled a large compendium of small RNA datasets in the extracellular (EC) or the intracellular (IC) compartment. These datasets, along with their ‘EC’ versus ‘IC’ labels, were obtained from three distinct sources: (i) a dataset of intracellular and extracellular small RNAs (between 18 and 50nt) we had previously generated across eight cell line models^7^, (ii) the Extracellular RNA Communication Consortium Atlas (exRNA Atlas)^13^ dataset, and (iii) The Cancer Genome Atlas (TCGA) small RNA sequencing data^14^. Given that the cell-free RNA content is not correlated with the abundance of small RNAs in the cell, we hypothesized that a *cis*-regulatory grammar serves as a localization signal for small RNA sorting into extracellular space. First, to search for EC-associated RNA sequence and structural features, we compiled the primary sequence, k-mer frequencies (k=1,2,3,4), *in silico* folding free energy, and predicted secondary structures as input features to train our model (**Fig 1A**). Starting with simpler models, we trained linear SVM, Gaussian kernel SVM, and random forests as classifiers. The poor performance of these models motivated us to train more complex models with increased learning capacity. We tested various neural network architectures, starting with shallow convolutional neural networks (CNN) and recurrent neural networks (RNN), as well as DeepBind, a previously developed CNN model ^15^. Upon hyperparameter tuning, we observed an increase in performance upon switching to a Gated recurrent units (GRUs)-based deep recurrent neural network architecture (**Fig 1B**). As shown in **Fig S1A**, we benchmarked our GRU model, which we named “ExoGRU’’ (**Figure 1C**), against several existing machine learning and deep learning models. **Figure 1D** shows the performance of ExoGRU, evaluated on the held-out test set, in which we achieved an area under receiver operating characteristic (AUROC) of 0.95 and an area under the precision recall curve (AUPRC) of 0.8, respectively. At 83% specificity, the sensitivity of ExoGRU was 91% (see the confusion matrix in **Fig S1B**). We also sought to assess the contribution of each input feature to the performance of ExoGRU. From our initial list of features described above, we observed that the primary sequence alone is sufficient to effectively distinguish IC sequences from EC.

**Figure 1.**
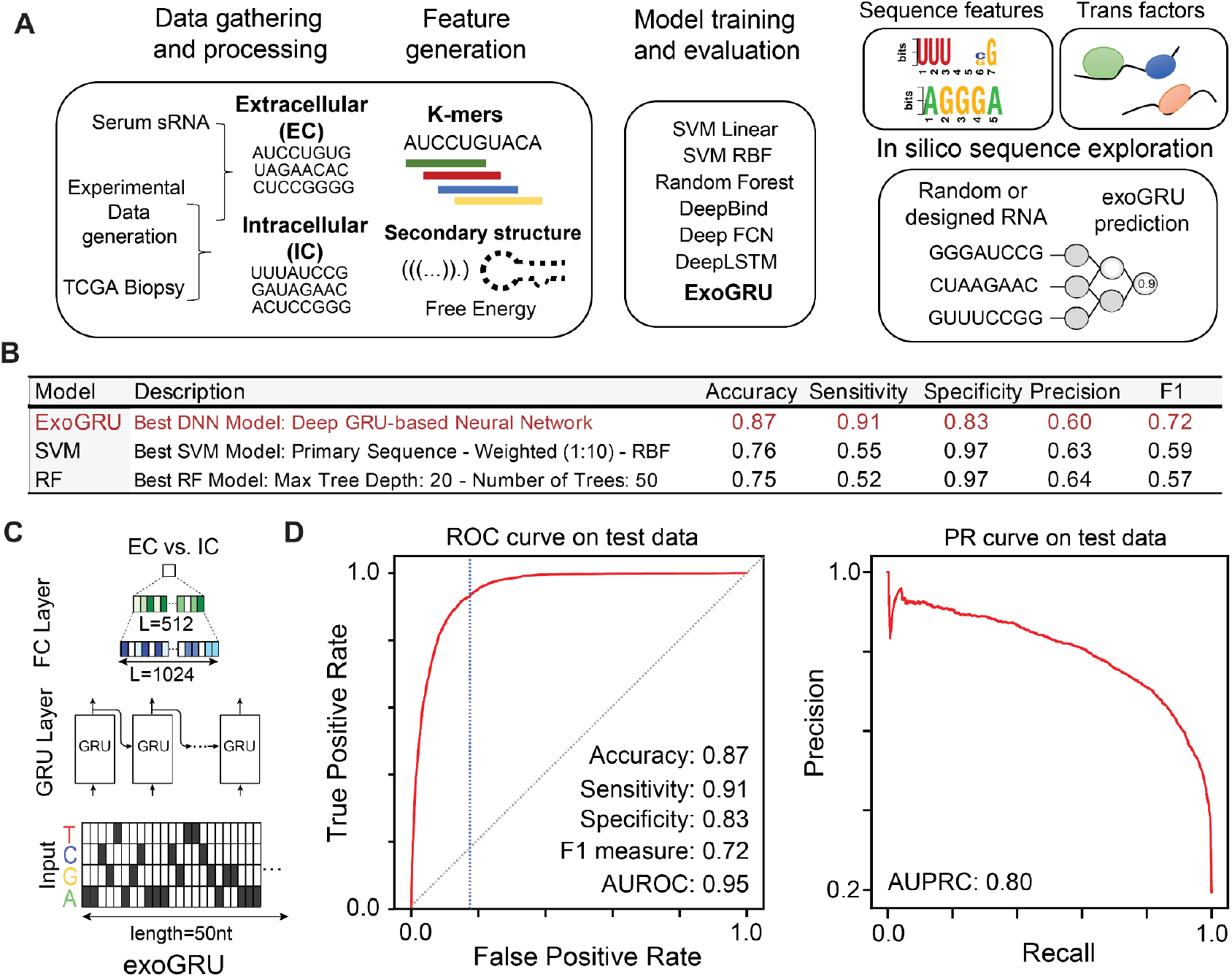
Predicting small RNA secretion from RNA sequence and structural features. **(A-B)** An overview of our strategy in this study: we used in-house and publicly available data to curate a dataset of intracellular and cell-free small RNA species. Following extensive feature engineering and evaluating various modeling strategies, we selected the best machine learning models for prediction of small RNA secretion. We observed that ExoGRU, a recurrent neural network model, outperforms other models in this task. We then performed feature attribution scoring and model dissection to dissect the cis-regualtory grammar captured by ExoGRU. **(C)** The architecture of ExoGRU following hyperparameter optimization. **(D)** Receiver Operating Characteristic (ROC) and Precision-Recall (PR) curves for the ExoGRU model for the held-out test set. Positive samples are the extracellular (EC) sequences and negative samples are the intracellular (IC) ones. The performance metrics of this model are also listed.

### Experimental verification of ExoGRU predictions

To further evaluate the performance of our model, we sought to focus on small RNAs whose status is predicted by ExoGRU with high confidence, i.e., focusing on high confidence true-positives and negatives. For this, we used ExoGRU to select those sequences with the highest and lowest secretion probabilities and labeled them as ECX (high confidence EC) and ICX (high confidence IC) (**Fig S1C-D**). We next implemented a variety of approaches to experimentally verify the ability of ExoGRU to capture the small RNA secretory grammar among these sequences. First, we generated an exogenously expressed small RNA library composed of two different sets of sequences: high confidence secreted small RNAs (ECX), and mutated variants of ECX (MUT). The latter set of sequences was generated by randomly mutating ECX small RNAs, in one or two positions, so that ExoGRU no longer classified them as secreted RNAs. We cloned this library, containing both ECX and MUT sequences, in a lentiviral construct downstream of a U6 promoter (pLKO.1 backbone) ^16^. We then transduced the MDA-MB-231 breast cancer cell line, which was among the lines used in our original dataset ^7,13,14^. We isolated small RNAs from extracellular vesicles (EV), conditioned media (CM) and intracellular (IC) fractions of this library and performed small RNA sequencing across all samples in biological replicates. We then aligned the resulting reads to the reference library to assess the abundance of each small RNA in the extracellular and intracellular space. It should be noted that expressing small RNAs via an exogenous construct may result in RNA species that (i) are mis-localized and therefore rapidly degraded, and (ii) lack the endogenous molecular context they rely on for successful secretion. Of the 600 pairs tested, in 55 cases, both the ECX and MUT pairs were stably expressed and therefore successfully captured by our assay. In order to assign a secretion score to each small RNA, we compared its abundance in the extracellular fractions (EV or CM) to intracellular RNA (IC). We observed that the resulting ‘enrichment scores’ were significantly higher for extracellular small RNAs (ECX) compared to their mutated counterparts (MUT), in both CM and EV fractions (**Fig 2A**). We observed a large fraction (93%) of exogenously expressed ECX sequences were indeed secreted; and more importantly, slight modifications to these sequences, guided by ExoGRU, resulted in a substantial and significant drop in their secretion potential. To assess the concordance between experimental measurements and ExoGRU predictions, we used a ROC curve to measure the association between experimental and ExoGRU labels at every classification threshold across the CM enrichment score (**Fig 2B**). We used the threshold resulting in a specificity of 0.75 to make “EC” and “IC” calls based on the experimental CM enrichment score. We used the resulting experimental classes to generate a confusion matrix against the ExoGRU labels and calculate performance metrics (**Fig 2C**). We also performed a similar analysis for the EV fraction, presented in **Fig S2B-C**, by calculating an experimental EV-enrichment score. Our observations in the EV fraction were similar to the conditioned media, albeit with a lower performance (70% accuracy versus 82% in CM). This was not unexpected since EV purification often suffers from technical variation and the recovered RNA levels are substantially lower.

**Figure 2.).**
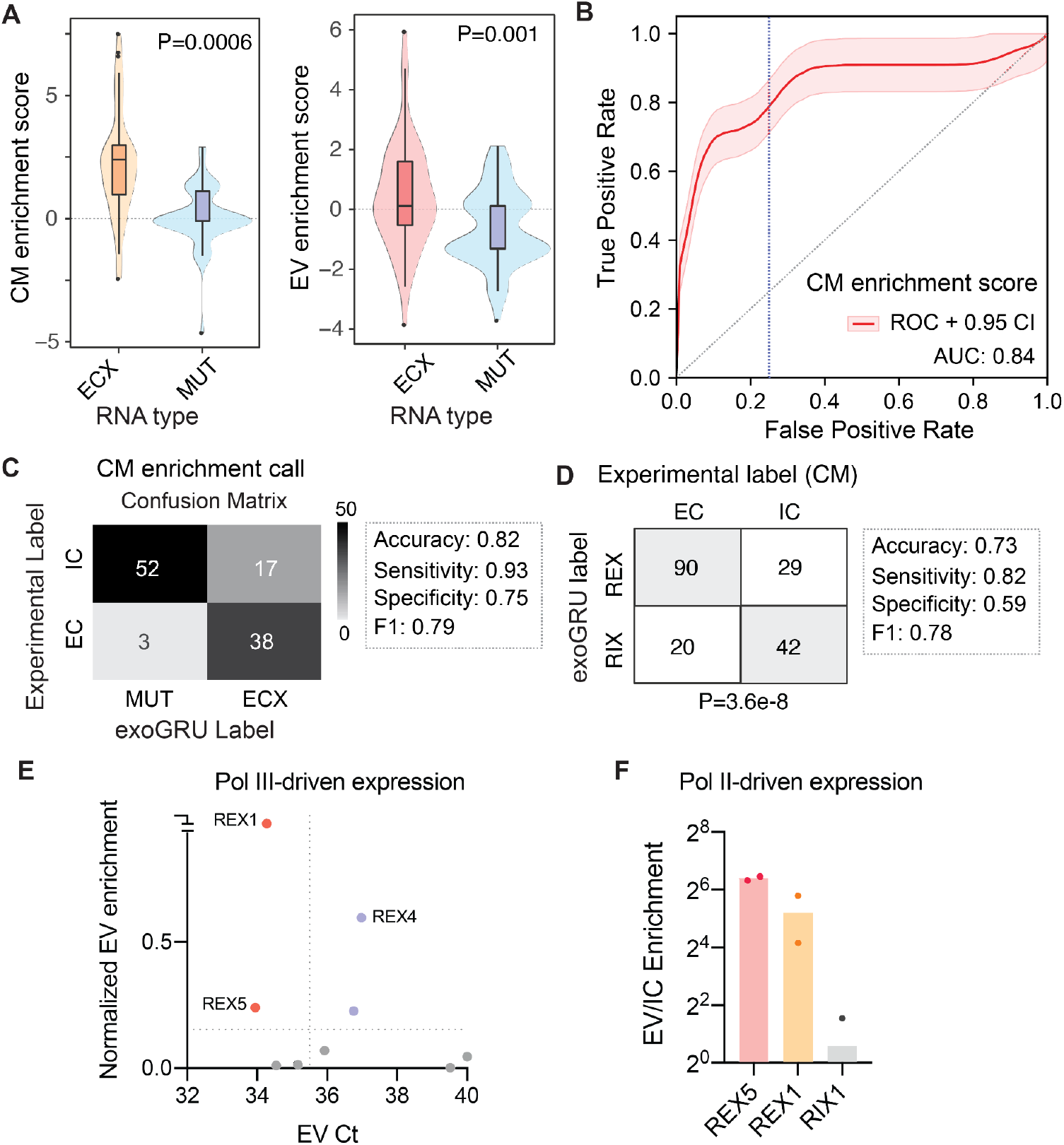
Experimental validations of ExoGRU predictions. **A)** EVX vs muted EVX smRNA distribution in EV or CM fractions. **B)** ROC curve for ExoGRU predicted EVX sequence enriched in CM. The smoothened ROC curve was generated by performing 1000 bootstraps. **C)** Confusion matrix for EVX and mutated EVX exoCNN labels and their experimentally validated EV vs IC distribution. Samples taken from CM and IC. **D)** ExoCNN predicted REX and RIX sequences and their experimentally validated EC vs IC distribution. **E)** Ct values of REX1-8 and RIX1-3 sequences in EV. All sequences were cloned under RNA polymerase III promoter. **F)** EV enrichment of REX1, REX5 and RIX1 sequences validated by qPCR. Sequences were cloned under RNA polymerase II promoter.

### The ability of ExoGRU to generalize its predictions to synthetic sequences

We next sought to determine whether ExoGRU can be used for generation, as opposed to mere classification, of synthetic small RNA sequences that are secreted effectively. Furthermore, we sought to assess whether the patterns learned by ExoGRU based on natural small RNAs is sufficiently generalizable to predict secretion probability of synthetic sequences. To these ends, we randomly generated RNA sequences with an average length of 20nt and dinucleotide frequencies matching those observed in the endogenous small RNAs. We then used ExoGRU to estimate their probability of secretion and selected ~400 sequences that were classified as ‘EC’ (labeled REX for randomly generated high confidence EC) and a similar number that were classified as ‘IC’ (labeled RIX for randomly generated high confidence IC). We synthesized REX/RIX sequences and cloned them similar to the above. Finally, we transduced this library into MDA-MB-231 cells and profiled small RNAs from the conditioned media (CM) and extracellular vesicles (EV). If a given randomly generated sequence was observed in the extracellular fraction, it was given the ‘EC’ label, otherwise, it was labeled as IC. In **Fig 2D**, we have provided the resulting contingency table comparing the experimental and computational labels. The accuracy of ExoGRU class predictions for these synthetic sequences was 73%, with 82% sensitivity and 59% specificity. We also used a χ2 test to calculate a *p*-value for the observed counts (P=3.6e-8).

We were intrigued by the ability of ExoGRU to generalize well to previously unseen sequences and to effectively identify entirely synthetic sequences that are efficiently secreted. Therefore, to independently verify the patterns observed for the REX and RIX sets in our sequencing data, we selected eight REX sequences (REX1 through REX8) and three RIX sequences (RIX1 through RIX3). We cloned these under a Pol III promoter (the same pLKO.1 backbone as the library) and generated MDA-MB-231 cell lines for each construct individually. After isolating small RNAs from EV and IC fractions in biological replicates, we performed quantitative RT-PCR to compare the enrichment of each sequence in the extracellular fraction. We used both the abundance and enrichment of small RNAs in the EV fraction as our selection criteria. We required the Ct values in EV fraction to be below 35 and log-fold EV enrichment to be above 0.1. We used miR-16, which is abundantly secreted, as an endogenous control in this assay and both IC and EV values were first normalized to miR-16. As shown in **Fig 2E**, REX1 and REX5 were significantly enriched in the EV fraction and were validated as bonafide EV-associated small RNAs with both small RNA sequencing and targeted RT-qPCR. Finally, we also tested the expression and secretion of REX1 and REX5 in MDA-MB-231 cells when cloned under a CMV promoter in the BdLV backbone^17^. To do so we used self-cleaving ribozymes^18^ to express our REX/RIX sequences under this Pol II promoter. In this case as well, we observed a significant enrichment of REX1 and REX5 in the EV fraction (**Fig 2F**). Together, our results validate the performance and utility of ExoGRU both as a predictive model that captures the small RNA secretory grammar and also as a generative model that can nominate synthetic small RNAs that are effectively secreted.

### Gaining insights into the RNA secretory mechanisms by dissecting the grammar learned by ExoGRU

ExoGRU effectively captures the probability of secretion from the primary RNA sequence alone, which implies the presence of an underlying shared sequence grammar that governs this process. *Cis*-regulatory elements often mediate interactions with master regulators, such as RNA binding protein, to influence the RNA life cycle. In fact, several RNA binding proteins have already been shown to play a direct role in RNA sorting into exosomes^1^. In order to systematically explore the role of RBPs in small RNA sorting and secretion, we first focused on applying motif discovery methods to the ECX and ICX sequences to find highly discriminative and class-specific motifs. We used three separate motif finding strategies, namely MEME^19^, Homer^20^, and FIRE^21^. We identified multiple sequence motifs that were enriched specifically in the ECX sequences. In parallel, we also used CLIP-seq data from the RNA ENCODE project^22^ to identify RBPs whose binding sites are enriched among the secreted RNAs. Using signal and peak-calling results of each RBP, and genome coordinates of ECX and ICX sequences, we sought to identify RBPs that are enriched for interactions with the ECX sequences. We applied the Mann–Whitney statistical test to detect such significantly greater overall signal values among the ECX and ICX regions. By intersecting these motif and RBP analyses, we arrived at six potential RNA-binding proteins that could preferentially bind to secreted small RNAs (**Fig S3A**).

Of these, we focused on YBX1, HNRNPA2B1, and RBM24 since they are also found within the extracellular space ^23^. As shown in **Fig 3A**, the known motifs for these RBPs were significantly enriched among cell-free small RNAs, even when controlled for length and dinucleotide content. Re-identification of YBX1 through this approach serves as a validation of our strategy given that it is known to be a major factor in microRNA and small RNA sorting into the exosomal compartment ^24^. Similarly, while not as well-characterized, HNRNPA2B1 has also been previously implicated in microRNA sorting ^25^. RBM24, on the other hand, does not have a canonical role in RNA secretion; however, it is known to be present within exosomes ^26^. To further explore the role of HNRNPA2B1 and RBM24 in small RNA secretion, we used CRISPR-interference to knockdown these RBPs and measure their consequences on the cell-free RNA content. We achieved a 77% knockdown for HNRNPA2B1 and 88% for RBM24 in MDA-MB-231 cells using lentiviral transduction as described in methods. We then isolated RNA from EV, CM and IC compartments for small RNA sequencing. As shown in **Fig 3B-C**, silencing HNRNPA2B1 and RBM24 resulted in a significant reduction in the abundance of small RNAs that contained their binding sites in both the extracellular vesicles (EV) and conditioned media (CM) fractions. This observation confirms the involvement of these RNA-binding proteins in RNA sorting and secretion.

**Figure 3.**
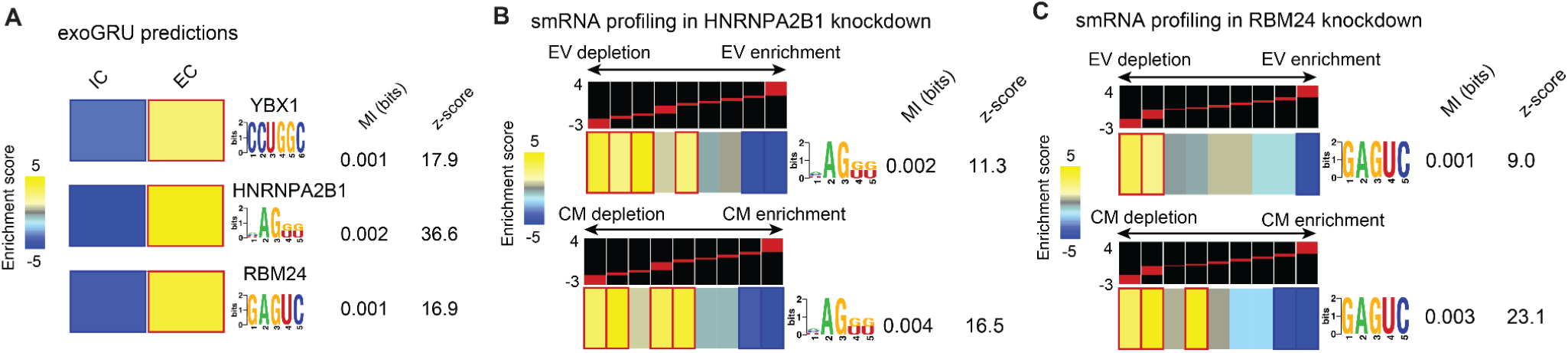
Use of ExoGRU in dissecting RNA secretory mechanisms. **A)** YBX1, HNRNPA2B1 and RBM24 motifs as predicted by exoGRU to be enriched in EC. **B)** Heatmap showing enrichment score of smRNAs containing HNRNPA2B1 motifs in EV upon decreasing HNRNPA2B1 expression. **C)** Heatmap showing enrichment score of smRNAs containing RBM24 motifs in EV upon decreasing RBM24 expression.

We next sought to confirm that, as previously claimed, HNRNPA2B1 sorts small RNAs it binds into exosomes. For this, we took advantage of UV-crosslinking co-immunoprecipitation followed by sequencing. CLIP-seq often includes a nuclease digestion step to footprint RBP binding sites across the transcriptome; however, by omitting this step, the bonafide small RNA targets bound by an RBP of interest can be profiled instead. We have previously used this approach for other RNA-binding proteins, such as AGO2 and YBX1 ^27^. Visualization of radiolabeled RNA crosslinked to HNRNPA2B1 on a denaturing gel revealed a faint but visible band at the correct size range (**Fig S3B**). We extracted these HNRNPA2B1-bound RNAs and performed high-throughput sequencing. Motif analysis of the identified binding site showed a strong and highly significant enrichment of the HNRNPA2B1 motif among the bound small RNAs (**Fig S3C**), which serves as a technical quality control. Finally, we asked whether these HNRNPA2B1-bound small RNAs were among those depleted from the exosomal space upon HNRNPA2B1 knockdown. Consistently, we observed a marked reduction in the secretion of these RNA, with a higher statistical significance compared to the HNRNPA2B1 motif analysis (**Fig S3D**). Together with the prior reports, our results show that HNRNPA2B1 binding to small RNAs is required for their effective secretion.

### HNRNPA2B1 and RBM24 exoCLIP shows enrichment of EV predicted sequences

The presence of RNA-binding proteins HNRNPA2B1 and RBM24 in extracellular vesicles along with their putative small RNA targets strongly suggests direct interactions within the exosomal space. However, direct evidence of RNA binding and the identity of their target RNAs remained lacking. To tackle this problem, we developed a novel approach for capturing the specific RNA molecules that a given RBP interacts with in the exosomal space. This approach, which we have named exoCLIP, is similar to CLIP-seq but uses UV treatment of conditioned media to crosslink RBP-RNA complexes in the cell-free fraction (**Fig 4A**). Using exoCLIP, we sought to demonstrate a direct interaction between HNRNPA2B1 and RBM24 and their target small RNAs. In the case of HNRNPA2B1, we tested both the A2 and B1 isoforms. We transduced MDA-MB-231 cells with FLAG-tagged copies of HNRNPA2, HNRNPB1, and RBM24, respectively. We then performed exoCLIP-seq for each line using FLAG co-immunoprecipitation. We used CLIP Toolkit ^28^ to call peaks for each of the RBPs using two strategies; one based on sequence coverage or signal, and the other based on crosslinking induced mutations (CIMs). Both strategies yielded between hundreds and thousands of RNA targets, a fraction of which mapped to annotated small RNAs (**Fig S4A**). These results indicate that HNRNPA2B1 and RBM24 indeed bind their RNA targets directly in the cell-free space. Interestingly, while we observed some correlation between the HNRNPA2 and HNRNPB1 isoforms, there were also many isoform specific binding sites for these RBPs (**Fig S4B**). In **Fig 4C**, we have included examples of small RNAs, in this case tRNA fragments, that are bound by each RBP as evidenced by the exoCLIP signal and the presence of crosslinking induced deletions (CIDs).

**Figure 4.**
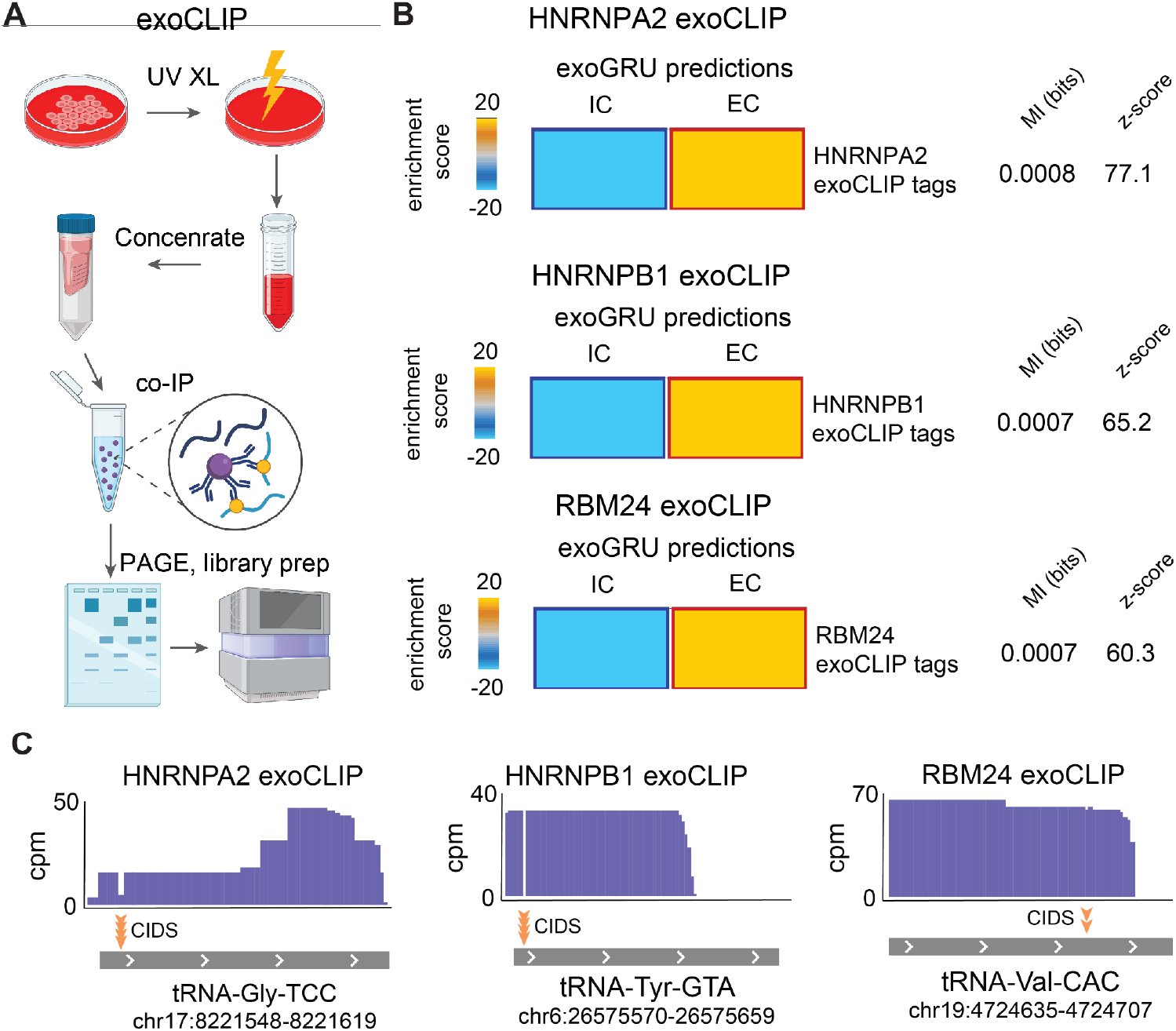
Applying exoCLIP to look at the enrichment of HNRNPA2B1 and RBM24 bound smRNA sequences in cell free media. **A)** overview of exoCLIP workflow: UV treatment of conditioned media to cross link RBP-RNA complexes and using co-IP to pull down the RBP-RNA complexes of interest followed by RNA library preparation and sequencing. **B)** Overlay of HNRNPA2, HNRNPB1 and RBM24 exoCLIP results on ExoGRU EC and IC bound smRNA predictions and looking at their enrichment scores. **C)** Examples of smRNAs bound by HNRNPA2, HNRNPB1 and RBM24 extracted from the exoCLIP sequencing results and their CIDs sites.

Since we had selected HNRNPA2B1 and RBM24 based on our analysis of high-confidence predictions for EC and IC RNAs from ExoGRU, we expected these predictions to match the exoCLIP results as well. To assess this possibility, we measured the enrichment of bound small RNAs from each dataset among the EC vs IC small RNAs. As shown in **Fig 4B**, we observed a significant over-representation of EC small RNAs that are directly bound by HNRNPA2B1 and RBM24.

## Discussion

It is hypothesized that extracellular small RNAs play a key role in intercellular communications and regulation of various biological processes ^1,5,6,29,30^. Identifying these specific RNA molecules and understanding their mechanisms of action has led to the discovery of different disease associated biomarkers and therapeutic targets ^7–9,29,31–33^. However, our understanding of how these RNA molecules are sorted and delivered into the extracellular space is still limited.

Multiple studies have identified different RBPs responsible for RNA secretion into the extracellular space ^24,25^. However, the full mechanisms underlying smRNA delivery are still largely unknown. A recent study comparing intracellular vs. extracellular miRNA profiles found multiple “EXOmotifs” and “CELLmotifs” on miRNA responsible for their secretion from or retention in the cells, suggesting that there are various different motifs and RBPs involved in this process ^10^. While the study provided valuable information on miRNA distribution in metabolically important cells, we aimed to further explore the mechanisms behind small RNA sorting in cancer cells using machine learning tools and novel molecular biology approaches.

To further decipher the principles of small RNA delivery to the extracellular space, we asked three specific questions: (1) which RNA sequences are selected and secreted; (2) can we develop a computational model that learns the sequence grammar that underlies RNA secretion; and (3) using this model, can we learn the molecular mechanisms that drive this selection? To tackle these questions, we developed ExoGRU, a deep recurrent neural network, to predict the secretion probability for any small RNA given the primary sequence. We rigorously verified our model’s ability to capture the small RNA secretory grammar by testing the impact of ExoGRU-guided targeted mutations on the secretion of endogenous small RNAs. We found the RNA primary sequence to be sufficient to discriminate between the intra- and extracellular small RNAs.

Additionally, we used exoGRU to reveal the regulatory grammar captured by the model. Using motif discovery methods and CLIP-seq data combined with high-confidence ExoGRU predictions, we identified several RBPs that preferentially bind to secreted RNAs and are associated with the RNA sorting process. In addition to recapitulating the known involvement of YBX1, we also demonstrated the role of RBM24 and HNRNPA2B1 in RNA secretion through CRISPR-interference and CLIP-seq. We described exoCLIP, a novel approach to capture direct RBP-RNA interactions in cell free media. Using this method, we successfully characterized RBM24, HNRNPA2 and HNRNPB1 RNA targets in the extracellular space. Our exoCLIP-seq data also aligned with ExoGRU predictions as we saw enrichment of EC associated sequences in these data. Overall, our results demonstrate the performance and utility of ExoGRU as a predictive model that captures the small RNA secretory grammar, and provides insights into the role of RBPs in small RNA sorting and secretion.

Last but not least, we showed ExoGRU’s prediction ability is generalizable to synthetic sequences. This was demonstrated through sequencing and quantitative PCR analysis of randomly-generated but ExoGRU-scored libraries of EC and IC sequences (REX/RIX). The validation process further confirmed the accuracy of the predictions made by the ExoGRU model. Using this feature of ExoGRU we will be able to design fully engineered and efficiently secreted sequences that can be used as biomarkers as well as having further applications in synthetic biology.

## Acknowledgements

We thank Dr. Sohail Tavazoie for his support and his comments on this manuscript. We also thank Drs. Babak Alipanahi and Hamed Najafabadi for his feedback on the earlier versions of this study. Lastly, we would like to acknowledge the UCSF Center for Advanced Technology (CAT) for high throughput sequencing and other genomic analyses.

## Code Availability

The code used in this study is available on GitHub at https://github.com/Exosomians/exosomians. The code was developed using the Python and R programming language. The code was designed to reproduce the analyses presented in this manuscript and may be useful for researchers wishing to extend or replicate our findings. The GitHub repository includes documentation and instructions on how to use the code.

## Notes

### Competing Interest Statement

The authors have declared no competing interest.

### Summary of Updates

Author information

