## Supplementary Figures for "Revealing the Grammar of Small RNA Secretion Using Interpretable Machine Learning"

**A**

| model | hyperparameters | accuracy | sensitivity | specificity | precision | F1 |
|---|---|---|---|---|---|---|
| exoGRU | see Methods | 0.87 | 0.91 | 0.83 | 0.60 | 0.72 |
| exoNet | see Methods | 0.83 | 0.75 | 0.91 | 0.53 | 0.62 |
| exoLSTM | see Methods | 0.82 | 0.88 | 0.75 | 0.50 | 0.64 |
| DeepBind | see Methods | 0.79 | 0.81 | 0.77 | 0.47 | 0.59 |
| SVM | Just Primary Sequence - Weighted (1:10) - Linear Kernel | 0.79 | 0.70 | 0.88 | 0.33 | 0.45 |
| SVM | Weighted (1:10) - Linear Kernel | 0.79 | 0.70 | 0.87 | 0.32 | 0.44 |
| SVM | Just Primary Sequence - Weighted (1:10) - RBF Kernel | 0.76 | 0.55 | 0.97 | 0.63 | 0.59 |
| SVM | Weighted (1:10) - RBF Kernel | 0.76 | 0.54 | 0.97 | 0.60 | 0.57 |
| RF | Unweighted - Max Tree Depth: 20 - Number of Trees: 50 | 0.75 | 0.52 | 0.97 | 0.64 | 0.49 |
| RF | Weighted (Balanced) - Max Tree Depth: 20 - Number of Trees: 50 | 0.67 | 0.35 | 0.99 | 0.83 | 0.57 |
| RF | Weighted (Balanced) - Max Tree Depth: 20 - Number of Trees: 200 | 0.67 | 0.34 | 0.99 | 0.86 | 0.49 |
| SVM | Unweighted - Linear Kernel | 0.57 | 0.15 | 0.99 | 0.69 | 0.49 |

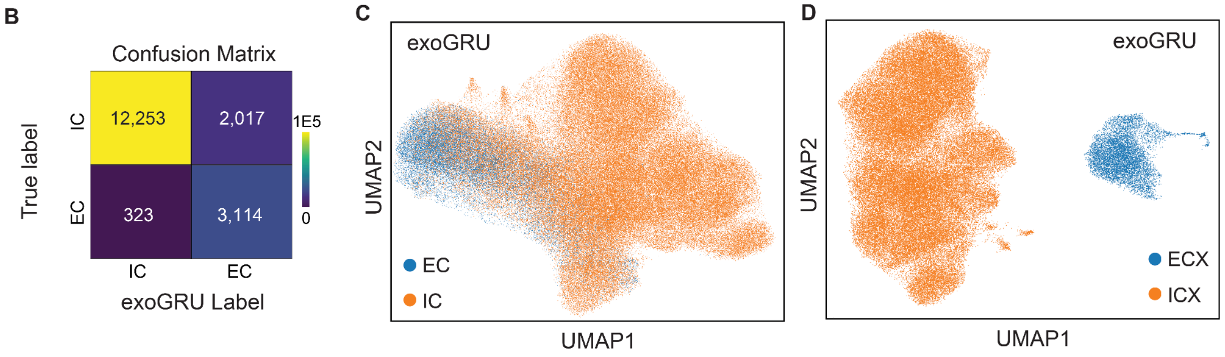


**Supplemental Figure 1. ExoGRU confusion matrix and embedding visualization. A)** Table compares accuracy, sensitivity,specificity and precision of different learning models tested **(B)** Confusion matrix for ExoGRU predictions for extracellular (EC) and intracellular (IC) labels. **(C)** UMAP projection was used to visualize the 64-dimensional embedding of EC vs IC. **D)** UMAP projection shows the 64-dimensional embedding of high confidence ECX and ICX calls.

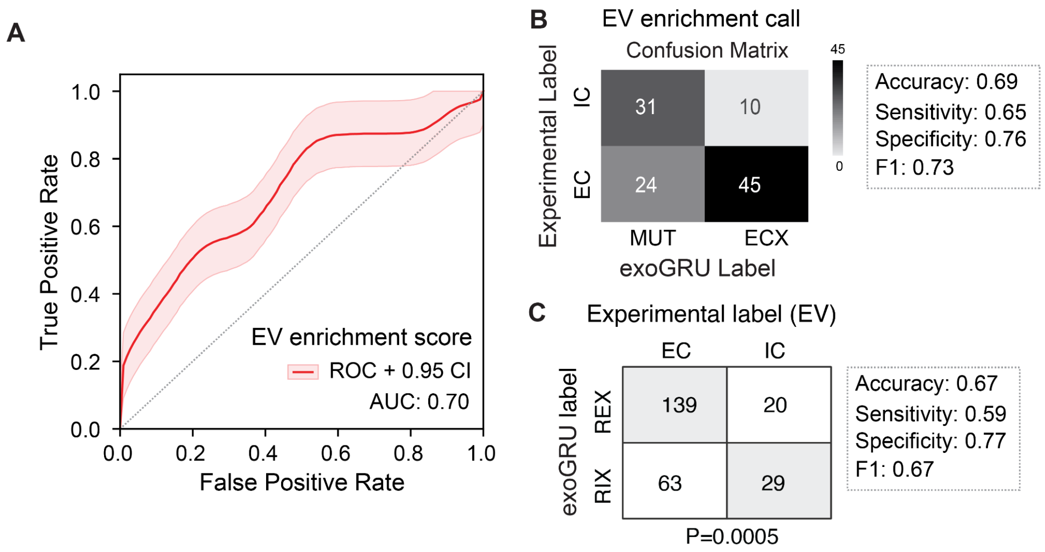


**Supplemental Figure 2. ROC curve and confusion matrix for ExoGRU predictions and its experimental validations A)** ROC curve for ExoGRU predicted EVX sequence enriched in CM. The smoothened ROC curve was generated by performing 1000 bootstraps. **B)** Confusion matrix for ECX and mutated ECX ExoGRU labels and their experimentally validated EC vs IC distribution. **C)** ExoGRU predicted REX and RIX sequences and their experimentally validated EC vs IC distribution.

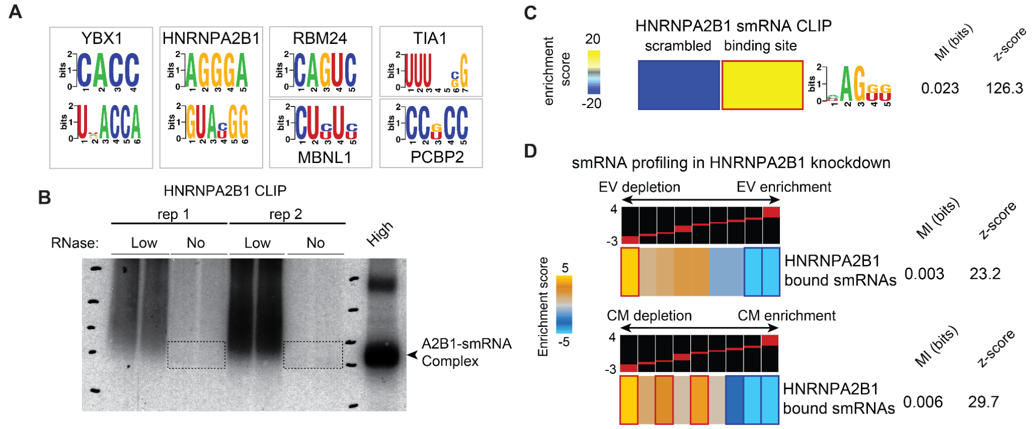


**Supplemental Figure 3. ExoGRU RBP motif discovery and CLIPseq to identify smRNA secretion mechanisms. A)** RNA binding protein motifs found by exoGRU to be enriched in ECX sequences and their corresponding RBPs. **B)** Western blot analysis of HNRNPA2B1 CLIP treated with zero, low and high RNase. **C)** motif analysis of HNRNPA2B1 CLIP sequencing result. **D)** Heatmap showing enrichment score of HNRNPA2B1 bound smRNA from CLIPseq data and their enrichment in EV and CM fractions upon HNRNPA2B1 knockdown.

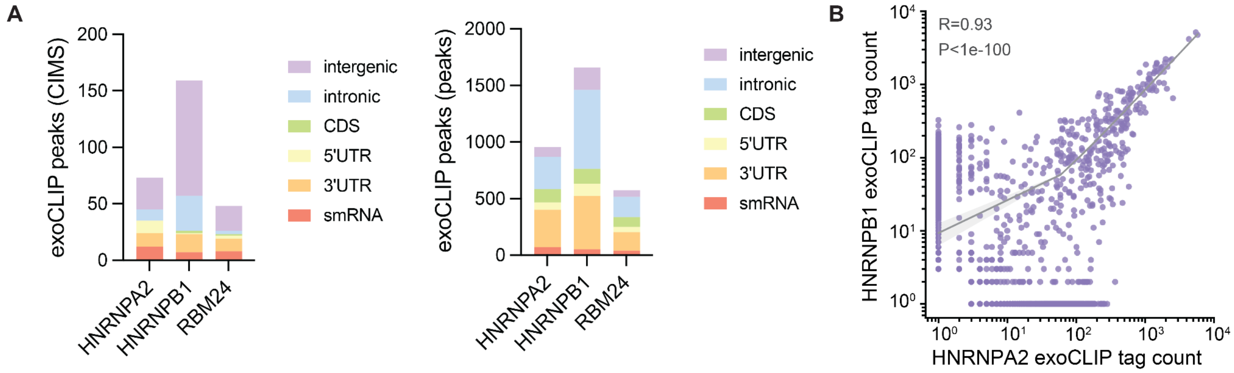


**Supplemental Figure 4. Annotated exoCLIP sequencing results A)** exoCLIP peaks of HNRNPA2B1 and RBM24 binding targets annotated to different RNA sites. **B)** Scatter plot shows the distribution of HNRNPA2 exoCLIP tags relative to HNRNPB1 tags.
