## Supplementary material for "Revealing the Grammar of Small RNA Secretion Using Interpretable Machine Learning": Materials and Methods

**Cell culture**

All cells were cultured in a 37°C 5% $CO_2$ humidified incubator. The MDA-MB-231 (ATCC HTB-26) breast cancer cell line, and 293T cells (ATCC CRL-3216) were cultured in DMEM high-glucose medium supplemented with 10% FBS, penicillin, streptomycin, and amphotericin B.

All the lentiviral constructs were co-transfected with pCMV-dR8.91 and pMD2.G plasmids using TransIT-Lenti (Mirus) into 293T cells, following manufacturer's protocol. Virus was harvested 48 hours post-transfection and passed through a 0.45 µm filter, and added to target cells 24 hours after they were seeded.

**Cell line generations:**

**MDA-MB-231 cells with RBP knockdowns**

Gene knockdowns were performed by first transducing MDA-MB-231 with dCas9-KRAB construct via lentiviral delivery of: pHR-UCOE-EF1a-dCas9-HAxNLS-XTEN80-KRAB-p2a-mCherry. MDA- dCas9-KRAB expressing cells were then sorted by FACS isolation of mCherry-positive cells. Guide RNA sequences for CRISPRi-mediated gene knockdown were cloned into pCRISPRia-v2 (Addgene #84832) [34] via BstXI-BlpI sites (see Supplementary table 1 for sgRNA sequences). After transduction with sgRNA lentivirus, MDA-MB-231 cells were selected with 2 µg/mL puromycin (Gibco). Knockdown of target genes was assessed by reverse transcription of total RNA to cDNA (Maxima H Minus RT, Thermo), then using sequence specific primers along with PerfeCTa SYBR Green SuperMix (QuantaBio) per the manufacturer's instruction. HPRT was used as an endogenous control (see Supplementary table 1 for  primer sequences).

**MDA-MB-231 cells overexpressing Flag-tagged RBPs**

For generation of flag tagged RBP cell lines, we cloned gblocks containing RBM24, HNRNPA2 or HNRNPB1 and the flag sequences into pLX302-EF1a plasmid via PacI-NheI sites.( Supplementary table 2 shows the gblock sequences). Plasmids were delivered to MDA-MB-231 by lentiviral transduction as described above. Expression of RBP-FLAG was assessed using western blot.

**MDA-MB-231 cells expressing REX or RIX sequences under pol III promoter**

For expressing REX1-8 / RIX1-3 sequences under U6 promoter we cloned oligos in Supplementary table 3 into pLKO.1 plasmid using AgeI and EcoRI sites, and transduced the MDA-MB-231 by lentiviral transduction as described above.

**MDA-MB-231 cells expressing REX or RIX sequences under pol II promoter**

For cloning REX1-3 and RIX1 sequences under the CMV promoter, we first cloned the ribozyme-small RNA-ribozyme (HH/HDV) cassette[18] into BdLV_Puro_mCherry using PacI and MluI site. We then digested the vector using AsiSI and cloned gblocks containing the sequence of interest (Supplementary table 4) using Gibson assembly.  Plasmids were delivered to MDA-MB-231 by lentiviral transduction as previously described.

**RT-qPCR for REX/RIX expression**

3.5 ul of isolated RNA was polyA tailed by adding 0.5ul 10X polyA polymerase buffer, 0.5ul 10mM ATP, 0.25ul polyA polymerase (NEB),0.25 ul H2O and incubating at 37°C for 10 minutes. 2.5 ul polyA tailed RNA was then reverse transcribed by adding 0.25ul 10mM dNTPs, 0.1ul 100uM dT T7 primer, 5X RT buffer,  0.15 ul RNAseOUT, 0.25 ul Maxima H Minus RT and 0.75 H2O by incubating at 50C for 15 minutes followed by 85C for 5 minutes. QPCR was done using PerfeCTa SYBR Green SuperMix, T7 primer, and miRNA specific primer as listed on Supplementary table 5. Mir16 primer was used as an endogenous control.

**Generation of smRNA libraries**

EVX/ICX and REX/RIX oligo pools were ordered from Twist Biosciences. Both oligo pools were separately cloned into pLKO.1-puro plasmid using AgeI and EcorI sites, and were transformed into MegaX electrocompetent cells with about 1000X coverage. The smRNA libraries were then transfected to MDA-MB-231 cells using lentivirus as described previously.

**RNA isolation from conditioned media (CM) and extracellular vesicles (EV)**

MDA cells were seeded in 10 cm or 15 cm plates. The next day the media was removed and cells were washed with 1X PBS. Cells were then incubated  in media prepared with exosome depleted FBS (cat# A2720801) in standard cell culture conditions for 48 hours. After 48 hours,

media was collected and spun down at 500g and passed through 0.4 um to remove any cells. For RNA isolation from conditioned media (CM), we took 1 ml of cell free media and performed RNA isolation using zymo research *Quick*-cfRNA Serum & Plasma Kit (cat# R1059).
The rest of the media was used for exosome isolation by adding Polyethylene Glycol 10000 (PEG, HR2-607) to 10 % final and overnight incubation at 4C. The next day PEG/media mixture was spun down at 3000g at 4C for 1 hour. We then removed the supernatant and proceeded to Zymo Research Quick-RNA Microprep Kit (cat#R1051) for RNA isolation from the EV pellets observed at the bottom of the tube.

**RNA isolation from cells**

Total RNA for RNA-seq and RT-qPCR was isolated using the Zymo Research Quick-RNA Microprep Kit (cat#R1051) with in-column DNase treatment per the manufacturer's protocol.

**ExoCLIP**

ExoCLIP of flag tagged RBM24, HNRNPA2, HNRNPB1 MDA-MB-231 cells was done by seeding 12M cells divided in four 15 cm cell culture plates for each cell line in DMEM media as described above. After 24 hours, the media was changed to DMEM with exosome free FBS. 48 hours after the media change, the conditioned media was collected and transferred to 50 ml falcon tubes and spun once at 500g and once at 2000g for 10 min at 4C to clarify the media from any cells. Clarified media was then transferred to 15 cm plates for crosslinking at 200 mJ/cm2 254 nm UV. After the first UV exposure we swirled the media and repeated the crosslinking step for a second time. Crosslinked clarified media was transferred to the centricon plus-70 filter 10K MWCO (millipore sigma UFC701008), and concentrated according to the manufacturer's protocol.
To the concentrated media we added protease inhibitor, SuperaseIN, EDTA, 1M Tris-HCl pH 7.5, and anti flag magnetic beads (CAT# A36797) and incubated with rotation for 20 hours at 4°C. Beads were magnetized and washed sequentially with cold low salt wash buffer, high salt wash buffer and PNK buffer two times each. This was followed by a PNK mediated dephosphorylation step (2.5ul 10X PNK buffer, 2ul 10X T4 PNK (10unit/ul), 0.5ul SuperaseIN, 20ul H2O) for 20 min at 37C and sequential washes with PNK buffer and high salt wash buffer. The de phosphorylated RNA-protein complexes were then poly A tailed using yeast PAP, PAP buffer, ATP and SuperaseIn (Jena 600U/ul) at 22 for 5 min. The poly A tailed RNA-Protein complex was then labeled by N3-dUTP, and yeast PAP, PAP buffer and SuperaseIn at 37C for 20 min. Beads were then washed by high salt wash buffer and PBS. The N3-labeled smRNA

was stained with 1mM 800cw DBCO at 22C for 30 min. Beads were magnetized and washed with high salt HITS-CLIP WB and PNK buffer respectively, and then resuspended in 20ul of 1X NuPAGE loading buffer + 50mM DTT final concentration diluted in PNK buffer, and heated at 75C for 10 min. Beads were placed on the magnet for elution. The eluted RNA protein complexes were then frozen in -80 and later used for WB analysis as described below.

**Low Salt Wash Buffer**

1X PBS (TC grade, no Mg++, no Ca++)

1% IGEPAL CA-630

**High Salt Wash Buffer**

5X PBS (TC grade, no Mg++, no Ca++)

1% IGEPAL CA-630

**1X PNK Buffer**

50mM Tris-Cl pH 7.4

10mM MgCl2

1% IGEPAL CA-630

**Western Blotting**

Eluted RNA-protein complexes from above were run on SDS-PAGE using 4-12% Bis-Tris NuPAGE gels, and transferred to protran BA-85 nitrocellulose membrane. The membrane was briefly rinsed in PBS, and placed in a sheet protector and imaged with a Licor Odessey instrument.

**Protein K digest and RNA capture**

The RNA-protein complexes imaged as described above appeared as a diffused signal with a modal size of ~15-20kDa above the expected MW of the protein of interest. Average MW of 21 nt long RNA is ~7kDa. Poly(A) tail ~20nt (~6.5kDa), therefore the position of the protein-RNA complex that will generate CLIP tags longer than 20nt is ~14kDa above the expected MW of the protein. HNRNPA2-Flag and HNRNPB1-Flag run at 38 and 39 kDa respectively and RBM24-Flag runs at ~28. Therefore, we cut between 55-85 kDa for HNRNPA2 and HNRNPB1 lanes and 39-70 kDa for RBM24 lane. The MDA only (no flag) lane was cut from 39-85 kDa.
The cut membranes were each transferred to a 1.5 ml Eppendorf tube and treated with 12.5 ul Proteinase K in 200 ul Proteinase K digestion buffer at 55C for 45 min. The samples were then

quickly spun down and the 200 ul of supernatant was transferred to a clean Eppendorf tube. Samples were then adjusted for salt by adding 19 ul 5M NaCl and 11 ul H2O per 200 ul sample. To capture the RNA we used 30 ul Oligo d(T)25 dynabeads (Invitrogen cat#61002) per IP. Beads were washed 2X with Proteinase K buffer before use. We transferred ~200ul salt-adjusted samples to the beads and incubated at 25C at 300 RPM for 20 min with occasional shaking of 1350 RPM. We then washed the samples/beads 2X with cold high salt wash buffer and 2X with PBS, magnetized and removed the supernatant. RNA was eluted by incubating the beads in 8 ul TE elution buffer at 50C for 5 min. Beads were magnetized and 7.5 ul of eluted RNA was transferred to clean PCR tubes.

**Small RNA library preparation**

Small RNA library preparation for samples taken from exoCLIP was done using Takara Bio SMARTer smRNA-Seq Kit (cat# 635029) with a few modifications. Since our RNA was already poly-A tailed, we skipped this step in the protocol and moved to the cDNA synthesis. We also wanted to incorporate UMI in our cDNA so we added 2.5ul smRNA mix 1 and 1ul of 10uM dT-UMI RT primer to our 7.5 ul poly A- tailed smRNA and incubated at 75C for 3 min and then placed on ice for 5 min. We then performed reverse transcription as described in the kit's protocol. In the PCR step we also added a 2 ul, 10 uM Universal reverse primer (P7) to the PCR mix and added the 78 ul mix to each cDNA sample. We then added the 2 ul index forward primer to each sample and incubated as described in the protocol. We purified the PCR product using Zymo Research select-a-size MagBead (cat#D4084-50).

Library preparation for RNA isolated from ECX/MUT or REX/RIX transduced MDA-MB-231 cells and small RNA library from HNRNPA2B1 KD and RBM24 KD MDA-MB-231 cells were prepared using an in house small RNA library preparation. 7.5 ul RNA was polyadenylated using 1ul NEB 10X polyA pol buffer, 1 ul 10 mM ATP, 0.25 ul RiboLock (40 u/ul), 0.25 ul E. coli PolyA pol. 5 u/µl (NEB), and incubated at 16C for 5 min, and then put on ice for maximum 5 min before proceeding to cDNA synthesis. We added 1 ul of RT primer, incubated at 72C for 3 min before putting on ice. We then prepared RT mix on ice using 2 ul 5X RT buffer with DTT (Thermo), 1 ul 10 mM dNTP, 4 ul 5M Betaine, 1 ul Maxima H- RT 200 u/µl (Thermo), 0.25 ul RiboLock 40 u/µl (Thermo), 1 ul 10 µM TSO-UMI primer. We incubated the RT mix at 42C for 30 min and then at 85C for 5 min. The cDNA amplification was carried by using 19 ul cDNA from previous step, 20 ul 5X Phusion HF buffer (Thermo), 2 ul 10 mM dNTP, 2 ul 12 µM Takara Fwd PCR primer, 2 ul 12 µM 12 µM Takara Rev PCR primer, 1 ul Phusion HS II pol. 2 u/µl (Thermo), and 48 ul H2O.

We then ran a PCR reaction for cDNA amplification as follows: 30 sec @ 98C - [10 sec @ 98C - 10 sec @ 65C - 5 sec @ 72C]xN cycles (N determined by performing a qPCR).

PCR reaction was purified through a MN NucleoSpin Gel & PCR Cleanup column (cat#740609), and eluted in 30 µl of water. We ran samples on a 8% TBE gel for 35 mins at 180V and stained the gel with 1X GelGreen in 1X TBE for 2 mins then imaged under Blue light. We cut the fragment of interest (150-200 bp) and placed the gel slices in a 0.5 mL tube with a hole pierced by a 18G needle. Spun the tube in a 1.5 mL tube until the gel was passed through the hole. We then added 400 µl of the DNA gel extraction buffer (10 mM Tris pH 8, 300 mM NaCl, 1 mM EDTA) to the gel, vortexed and froze on dry ice for 30 min, and then thawed overnight on a rotator. Next day we transferred the gel slurry to a Costar filter spin-column and spun at maximum speed until all liquid has passed through. We added 1.5 µl of GlycoBlue and 500 µl of isopropanol to the DNA solution, and put in -80C for 1 hour, then spun at 4C for 30 minutes, air dried for 10 minutes, and resuspended and incubated in 10 µl of 10 mM Tris pH 8 for 10 minutes.

**Sequencing**

Libraries were quantified using Qubit HS dsDNA kit, and also ran on an Agilent bioanalyzer HS DNA Chip or HSD1000 tapestation. All libraries were sequenced as SE65 runs on Illumina Hiseq 4000 at UCSF Center for Advanced Technologies.

**HITS-CLIP**

HITS-CLIP for endogenous HNRNPA2B1 was done as described by (Licatalosi et al., 2008) [35] with the modifications previously used for YBX1 small RNA CLIP (Goodarzi et al., 2015). MDA-MB-231 cells were UV-crosslinked at 400 $mJ/cm^2$ before cell lysis. Samples with and without RNase treatment were immunoprecipitated with an anti-HNRNPA2B1 antibody (Thermo, PA5-34939) for protein-RNA complexes. Polyphosphatase (Lucigen) was incubated with smRNA samples before ligation and PCR amplification with primers described by (Goodarzi et al., 2015). Constructed libraries were sequenced on the Illumina HiSeq2000 at the Rockefeller University Genomics Center.

**Data Acquisition**

To train our predictive models, we sourced small-RNA sequences data for intracellular and exosome-specific predictions from three reliable sources: Goodarzi et al. (GSE114366),

Extracellular RNA Communication Consortium Atlas (exRNA Atlas), and The Cancer Genome Atlas (TCGA). We first used the GSE114366 data, which was generated to investigate the roles of intracellular and extracellular small-RNAs in breast cancer. We extracted small-RNA-seq data of intracellular small-RNAs (IC) and small-RNAs present in extracellular vesicles (EV) from 8 different breast cancer cell lines. Although the small-RNA selection and secretion machinery may differ among different cell types and states, we assumed that there are some general and common mechanisms that exist in cells. Therefore, we merged all of the IC data, regardless of the cell line, and imported 30,093,690 IC-resided small-RNA sequences to our IC dataset. We also integrated all the EV data and collected 6,127,883 EV-resided small-RNA sequences to our EV dataset. Second, we imported 67,511,039 EV-resided small-RNA sequences from the exRNA Atlas, which were extracted from the serum part of blood cells of 12 samples. Third, we collected 1,8488,703 IC-resided miRNA sequences from the TCGA dataset which were extracted from normal cells of different tissues across the body. We selected miRNA-seq data and not RNA-seq data because our main focus in this research is to investigate the selection and secretion processes of small-RNAs inside a cell.

**Data Preprocessing**

After integrating data from three distinct sources, we performed several preprocessing steps to clean and organize the data. We first removed sequences that were present in both the IC and EV datasets, assuming that they belonged to the EV, as exported small-RNA sequences can also exist intracellularly. We then eliminated sequences that were less than 18 nts or more than 50 nts, as our focus was on small-RNAs. We also removed any sequences that contained "N" in their primary sequence in order to decrease ambiguity in our dataset. We eliminated duplicated sequences and sequences that were a substring of a bigger one. For example, if we had both ACGU and UACGU sequences in our dataset, we removed the former one. To further clean the data, we used the MEME-Suite's dust tool to mask and delete sequences that carried low-complex regions. The dust tool helps to identify and remove any non-informative regions in the sequences such as repetitive regions. After all of these preprocessing steps, we had 33,083 unique EV small-RNAs and 1,318,795 unique IC small-RNAs that were ready to be used for predictive model training.

### Feature Generation

Having datasets of IC and EV small-RNA primary sequences, we generated features to train our model. To generate secondary structures based on our primary sequences dataset, we used two secondary structure prediction tools: ViennaRNA and Forge. Using ViennaRNA, we were able to predict the dot-bracket form of the secondary structure and estimate the free energy of each small-RNA sequence. Additionally, we applied the Forge tool on the dot-bracket notation of sequences to predict the bulge-graph form of secondary structures. This way, for each small-RNA in our IC and EV dataset, we had three sequences an one quantity: primary sequence, dot-bracket secondary sequence, bulge-graph secondary sequence, and free energy. Subsequently, we extracted K-mers (K = 1,2,3,4) from the primary and secondary sequences mentioned above and calculated their frequency, which we used as input features for feature-based models. We normalized the features (K-mers frequency and free energy) based on the length of each sequence and also included the length of the sequence as a feature. Therefore, we generated different types of secondary structures and sequences from the primary ones to be used in sequence-based models, and then we extracted several features from the primary and secondary sequences to be utilized in feature-based models. Since 99.5% of our sequences' lengths were less than 50 bp,, we truncated sequences with length more than 50, and also padded all smaller sequences to have a fixed length. These preprocessings made our data consistent and ready to be fed into the model.

### Predictive Models

The predictive models examined in this research are divided into two categories: classical machine learning methods and deep learning methods. Classical methods trained/tested in this research include support vector machines (SVMs) and random forests (RFs). On the other hand, deep learning methods include models which are inspired by convolutional neural networks (CNNs) and recurrent neural networks (RNNs). It should be noted that due to the limitations of machine learning models (SVMs and RFs) in handling sequence-based data, all sequence-typed features were removed from the final design matrix for the machine learning experiments.

#### Support Vector Machines

Support Vector Machines (SVMs) are a family of supervised machine learning algorithms that are commonly used for linear and non-linear classification tasks, as well as for regression tasks. In this study, we conducted an ablation study to evaluate the effectiveness of different input feature spaces and kernel types for SVMs. Specifically, we defined six different training scenarios, which varied in terms of the input feature space and kernel type used.

To evaluate the effectiveness of non-Kmer features, we used two different feature sets: one that included all extracted features, and another that included only the K-mers extracted from the primary sequences. Additionally, we tested both linear and Radial Basis Function (RBF) kernels for the SVMs. Furthermore, due to the inherent class imbalance issue existing in the preprocessed dataset, we tried to mitigate this issue by using weighted SVM by weighting the class parameters inversely proportional to their sample frequencies.

**Random Forest**

In this study, we employed Random Forest, a powerful tree-based machine learning algorithm, to train on the preprocessed dataset. This choice was made due to the algorithm's ability to handle large feature sets and its robustness to overfitting compared to other existing machine learning models. Similar to the experiments conducted with Support Vector Machines (SVMs), we evaluated the effectiveness of both weighted and unweighted Random Forest models. Additionally, we investigated the impact of different tree population sizes on the performance of the algorithm. Specifically, we tested tree population sizes of 50 and 200 while keeping the tree-depth fixed at 20.

**ExoGRU**

ExoGRU, as the name suggests, consists of multiple GRU units stacked on top of each other. GRUs introduced a simpler alternative compared to LSTMs. They are able to capture relatively long-term dependencies by utilizing gates in order to control the information flow.

A GRU unit computes the hidden state at time step t as follows:

$$z_t = sigmoid(W_z x_t + U_z h_{t-1} + b_z)$$

$$r_t = sigmoid(W_r x_t + U_r h_{t-1} + b_r)$$

$$h'_t = tanh(W_h x_t + U_h(r_t * h_{t-1}) + b_h)$$

$$h_t = (1 - z_t)h_{t-1} + z_t h'_t$$

Where x_t is the input at time step t, h_{t-1} is the previous hidden state, W_z, U_z, W_r, U_r, W_h, U_h are the weight matrices, b_z, b_r, b_h are the bias terms and sigmoid and tanh are non-linear activation functions. The update gate z_t and reset gate r_t are used to control the flow of information into the hidden state h_t, allowing the network to better handle long-term dependencies.

**ExoLSTM**

ExoLSTM is also another network we employed in this study. The architecture consists of multiple LSTM units stacked on top of each other. Long Short-Term Memory (LSTM) units are a type of recurrent neural network (RNN) that uses a memory cell to store information over a longer period of time. The memory cell is controlled by gates that determine when to store, update, or discard information in the cell.

An LSTM unit computes the hidden state at time step t as follows:

$$i_t = sigmoid(W_i x_t + U_i h_{t-1} + b_i)$$

$$f_t = sigmoid(W_f x_t + U_f h_{t-1} + b_f)$$

$$o_t = sigmoid(W_o x_t + U_o h_{t-1} + b_o)$$

$$c_t = f_t * c_{t-1} + i_t * tanh(W_c x_t + U_c h_{t-1} + b_c)$$

$$h_t = o_t * tanh(c_t)$$

Where x_t is the input at time step t, h_{t-1} is the previous hidden state, c_{t-1} is the previous memory cell state, W_i, U_i, W_f, U_f, W_o, U_o, W_c, U_c are the weight matrices, b_i, b_f, b_o, b_c are the bias terms, and sigmoid and tanh are non-linear activation functions. The input gate i_t, forget gate f_t, output gate o_t and cell state c_t are used to control the flow of information into the hidden state h_t, allowing the network to better handle long-term dependencies.

**ExoCNN**

ExoCNN is a variant of convolutional neural networks (CNNs) designed to generate predictions from sequences. The architecture of ExoCNN is composed of several layers, including convolution, pooling and fully connected layers, each of which contains tunable weights and biases. One key aspect of the ExoCNN architecture is the use of "conv blocks" as firstly defined in VGG[36] which are composed of multiple consecutive convolution layers followed by a max-pooling operation. In the max-pooling operation, the maximum value is computed for each window of size 2 in the "conv block"'s output matrix, this helps to summarize spatial information in to the output while retaining the conserving the spatial information. Following the "conv blocks" and max-pooling operations, the output of the last max-pooling operation is flattened and fed to a classifier head with 2-layer fully connected neural network. Similar to the convolution layers, rectified linear activation functions are used in the head. The number of neurons in the hidden layers of ExoCNN's classification head are 1024 and 128 respectively. Finally, the output of the last layer is passed through a sigmoid function which generates a (secretion) probability for the input sequence.

**Model Training**

As shown in Table 1, the sample frequency of the IC class is significantly higher than the EV class, which results in a common problem in machine learning known as class imbalance. To address this issue, we downsampled the IC dataset to balance the class frequencies. Additionally, we employed the weighted cross-entropy (WCCE) loss function for training our ExoCNN model in which each class weight is inversely proportional to its sample frequency. The original EV dataset and the downsampled IC dataset were used to train our predictive models. To assess the performance of deep learning models, we performed stratified train/validation/test split with proportions of 0.8, 0.1, and 0.1 on our preprocessed dataset.

We used the Adam optimizer with a learning rate of 0.001 for 100 epochs for all DL models. We employed a batch size of 128 during training. To prevent overfitting, we employed early stopping and learning rate decay techniques during the training process. To initialize the weights of each layer in the network, we used the Xavier initializer. Additionally, we used L1 and L2 regularization techniques with a lambda value of 1e-6 to further prevent overfitting.

**Motif Discovery and Enrichment**

After training the network with small-RNA sequences, two sets of sequences were identified that the model was highly confident about being secreted or not. These sequences were named extreme extra-vesicular sequences (EVX) and extreme intra-cellular sequences (ICX), and had a confidence score above 0.95. In order to find motifs more accurately, we removed highly similar sequences from the EVX and ICX sets using the MEME-Suite's purge tool. To find the optimal similarity score threshold, we experimented with different thresholds and checked the number of sequences and removed ones for each threshold. Finally, we used a similarity score threshold of 50. The extreme sequences (EVX and ICX) were clustered based on edit distance and cosine distance. We found motifs based on both unclustered and clustered EVX and ICX, but the results were the same, so we eliminated the clustering step from the analysis pipeline. To perform an exhaustive motif search, we used several motif finders and tested various configurations of the tools. Three motif finding tools were able to discover motifs in the small-RNA sequences: MEME, Homer, and FIRE. We used these tools with three different input sets: EVX only, EVX vs ICX, and EVX vs randomly generated sequences that preserved di-nucleotide frequency. Using these three motif finding tools, three different configurations, and several parameter tuning, we found 10 motifs that were enriched in the EVX sequences. These motifs were related to previously known RNA-binding proteins that are involved in the secretion machinery, and were presented in Figure 4.

With the discovery of the EVX and ICX sequences and the corresponding motifs, we continued our research by identifying secretion-related RNA-binding proteins in two distinct ways. First, we compared the discovered motifs with already known ones in the literature and databases. This allowed us to identify any previously known motifs that were enriched in the EVX sequences and related to known RNA-binding proteins involved in the secretion machinery. Second, we analyzed the eCLIP-seq data of the ENCODE project to identify binding sites of human's RNA-binding proteins. This allowed us to identify any potential RNA-binding proteins that may be involved in the secretion of small-RNAs based on their binding sites in the EVX and ICX sequences.

**Motif Comparison**

We compared the 10 discovered motifs with known motifs of RNA-binding proteins to detect the proteins that are highly likely to bind to each motif and participate in the secretion machinery. To do this, we used three databases of Ray2013, RBPDB, and ATtRACT, and the MEME-Suite's Tomtom tool to find RNA-binding proteins (RBPs) that significantly bind to our discovered EV-enriched motifs. This motif comparison process gave us 7 proteins that are highly likely to bind to our secretion-related motifs, as shown in Figure 4. As previously mentioned, two of these proteins have already been verified to be involved in the secretion machinery. This comparison process helps us to identify potential players in the secretion process and further investigate them.

**RBP Binding Sites Analysis**

We aimed to identify RNA binding proteins (RBPs) in the ENCODE database that may have greater interactions with extreme EV sequences, as opposed to IC sequences. Our hypothesis is that these proteins may play a role in the secretion machinery. We filtered out proteins that did not have signals (bigWig file) or peaks (BED file) as their output type and that were not based on the GRCh38 reference genome. This resulted in a final selection of approximately 150 proteins.

To begin, we determined the maximum signal value at each nucleotide position for a specific protein if we have multiple experiments (bigWig files). Next, we extracted signal values for nucleotide positions that overlapped with peak regions, separately for IC and extreme EV sequences. We then used the Mann-Whitney statistical test to compare these two sets of signal values and calculate a p-value to determine if the EV signals were significantly greater than the IC ones.

To obtain comparable signal intensity values and gain a deeper understanding of the interactions between EV extreme sequences and RBPs, we evaluated various scoring methods. In our initial analysis, we obtained signal values (scores) for EV sequences and assigned zero values to regions that did not overlap with peak regions for a specific protein. We also applied this method to IC sequences. This resulted in many zero values in our scores, and the Mann-Whitney test showed a significant sensitivity to the mean in these scenarios.

To address this issue, we modified our approach. Instead of using all peak regions, we applied it to the union of IC and extreme EV regions. This eliminated many zero values from the scores and allowed us to better understand the natural behavior of RBPs. We found that they tend to bind to EV extreme sequences with high signal values and to IC extreme sequences with moderate signal values on average. We then used the Benjamini-Hochberg (BH) method to adjust our p-values and identified proteins with adjusted p-values less than 0.05 as being involved in the EV secretion machinery.

After analyzing the interactions between different proteins and RNA sequences, we also took an intra-protein approach to the problem. To obtain information within each sample (IC vs EV), we used extreme sequences of both IC and EV groups with a confidence probability greater than 0.9 that overlapped with peaked regions. We extracted several features including: the number of EV and IC extreme sequences overlapping with the peaked regions, the total length of each overlapping extremes with peaked regions, the total sum of the signal values for each overlapping extremes, and the mean value of signals for each of the extremes. With this data, we could assess the robustness and reliability of our results as a sanity check and also identify any potential outliers related to the secretion machinery. To do this, we used median absolute deviation (MAD), Z-test, and Percentile rank.
