## Supplementary Tables for "Revealing the Grammar of Small RNA Secretion Using Interpretable Machine Learning": Sheet1.html

|  | A | B | C |
| --- | --- | --- | --- |
| 1 | Table 1 |  |  |
| 2 |  | sgRNA sequence |  |
| 3 | sgRBM24 | ttgGCGGCTTCCGAAGTCGCCAGgtttaagagc |  |
| 4 | sgRBM24 | ttagctcttaaacCTGGCGACTTCGGAAGCCGCcaacaag |  |
| 5 | sgHNRNPA2B1\_1 | ttgGAGCAGCGCCGGGTCCCGTGgtttaagagc |  |
| 6 | sgHNRNPA2B1\_1 | ttagctcttaaacCACGGGACCCGGCGCTGCTCcaacaag |  |
| 7 |  | qPCR primers |  |
| 8 | RBM24\_q\_up | CAGAAAGGCCAACGTGAACC |  |
| 9 | RBM24\_q\_dw | TAGTGGGCAGGTATCCCGAA |  |
| 10 | HNRNPA2B1\_q\_up | GATATGGCAGTGGACGTGGAT |  |
| 11 | HNRNPA2B1\_q\_dw | CTCCATAACCGGGGCTACCT |  |
| 12 | HPRT-fwd | CAGTTAAAGTTGAGAGATCATCTCCACC |  |
| 13 | HPRT-rev | GCACTGAATAGAAATAGTGATAGATCCATTCC |  |
| 14 |  |  |  |
| 15 | Table 2 | Gblock sequence |  |
| 16 |  |  |  |
| 17 | Flag\_hnrnpa2 | aggaatttcgacatttaaatttaattaagctagcCACCATGgattacaaggatgacgacgataaaggcgcagactataaagacgatgatgacaagggttcaggatctggctccgggtcggagagagaaaaggaacagttccgtaagctctttattggtggcttaagctttgaaaccacagaagaaagtttaaggaactactacgaacaatggggaaagcttacagactgtgtggtaatgagggatcctgcaagcaaaagatcaagaggatttggttttgtaactttttcatccatggctgaggttgatgctgcaatggcagcaagacctcattcaattgatgggagagtagttgagccaaaacgtgctgtagcaagagaggaatctggaaaaccaggggctcatgtaactgtgaagaagctgtttgttggcggaattaaagaagatactgaggaacatcaccttagagattactttgaggaatatggaaagattgataccattgagataattactgataggcagtctggtaagaaaagagggtttgggtttgttacttttgatgaccatgatcctgtggataaaatcgtattgcagaaataccataccatcaatggtcataatgcagaagtaagaaaggctttgtctagacaagaaatgcaggaagttcagagttctaggagtggaagaggtgggaactttggctttggtgattcacgtgggggcggcggaaatttcggtccagggccaggcagtaactttagaggtggatctgatgggtacggtagtggacgtggatttggagatggctataatgggtatggagggggtcctggcgggggcaattttggagggagccccggttatggaggtggcagaggaggctatggcggtggaggacctggatatggcaaccagggtgggggctacggaggtggttatgacaactacggtggagggaattatggaagtggtaattacaatgattttggcaattataaccagcaaccttctaattacggtccaatgaagagtggaaactttggtggtagcaggaacatggggggaccatatggtggcggaaactatggtccaggaggaagtggaggtagtgggggttatggtgggaggagccgatactgatctcgacggt |  |
| 18 | Flag\_hnrnpb1 | aggaatttcgacatttaaatttaattaagctagcCACCATGgattacaaggatgacgacgataaaggcgcagactataaagacgatgatgacaagggttcaggatctggctccgggtcggagaaaactttagaaactgttcctttggagaggaaaaagagagaaaaggaacagttccgtaagctctttattggtggcttaagctttgaaaccacagaagaaagtttaaggaactactacgaacaatggggaaagcttacagactgtgtggtaatgagggatcctgcaagcaaaagatcaagaggatttggttttgtaactttttcatccatggctgaggttgatgctgcaatggcagcaagacctcattcaattgatgggagagtagttgagccaaaacgtgctgtagcaagagaggaatctggaaaaccaggggctcatgtaactgtgaagaagctgtttgttggcggaattaaagaagatactgaggaacatcaccttagagattactttgaggaatatggaaagattgataccattgagataattactgataggcagtctggtaagaaaagagggtttgggtttgttacttttgatgaccatgatcctgtggataaaatcgtattgcagaaataccataccatcaatggtcataatgcagaagtaagaaaggctttgtctagacaagaaatgcaggaagttcagagttctaggagtggaagaggtgggaactttggctttggtgattcacgtgggggcggcggaaatttcggtccagggccaggcagtaactttagaggtggatctgatgggtacggtagtggacgtggatttggagatggctataatgggtatggagggggtcctggcgggggcaattttggagggagccccggttatggaggtggcagaggaggctatggcggtggaggacctggatatggcaaccagggtgggggctacggaggtggttatgacaactacggtggagggaattatggaagtggtaattacaatgattttggcaattataaccagcaaccttctaattacggtccaatgaagagtggaaactttggtggtagcaggaacatggggggaccatatggtggcggaaactatggtccaggaggaagtggaggtagtgggggttatggtgggaggagccgatactgatctcgacggtatcggttaac |  |
| 19 | Flag\_rbm24 | aggaatttcgacatttaaatttaattaagctagcCACCATGgattacaaggatgacgacgataaaggcgcagactataaagacgatgatgacaagggttcaggatctggctccgggtcgcacacgacccagaaggacacgacgtacaccaagatcttcgtcggaggtctgccctatcacaccaccgacgccagcctgcgcaagtatttcgaggtcttcggagagatcgaggaggcggtggtcatcaccgaccggcagacgggcaagtcccggggctatggatttgtcaccatggctgaccgggctgctgccgaaagggcctgcaaggatcccaatcccatcattgatggcagaaaggccaacgtgaacctggcatacttaggagcaaaaccaaggatcatgcaaccaggttttgcctttggtgttcaacaacttcatccagcccttatacaaagacctttcgggatacctgcccactatgtctatccgcaggcttttgtgcagccgggagtggtcattccacacgtccagccgacagcagctgccgcctccaccaccccttacattgattacactggagctgcatacgcacaatactctgcagctgctgctgctgcagccgccgctgctgcctatgaccagtacccctatgcagcctctccagctgctgcaggatatgttactgctggaggttatggttatgcagtccagcagccaatcaccgcagctgcacctgggacagctgccgccgccgctgcagcagctgctgccgctgcagcatttggccagtaccagcctcagcaactgcaaacagaccgaatgcaata |  |
| 20 |  |  |  |
| 21 | Table 3 |  |  |
| 22 | REX\_1 | ggaaaggacgaaacaccggtTGCCCCTTCCTTCAGACAGCTTTTTTgaattctcgacctcgagaca |  |
| 23 | REX\_2 | ggaaaggacgaaacaccggtGTTGAGTGAAATCAGTTTCTTTTTTgaattctcgacctcgagaca |  |
| 24 | REX\_3 | ggaaaggacgaaacaccggtTCTCACTACTCTATTATTCTTTTTTgaattctcgacctcgagaca |  |
| 25 | REX\_4 | ggaaaggacgaaacaccggtACCCACGCCTAACTAATTCTTTTTTgaattctcgacctcgagaca |  |
| 26 | REX\_5 | ggaaaggacgaaacaccggtCAGTCCCACCTTCCCACTTTTTTTTgaattctcgacctcgagaca |  |
| 27 | REX\_6 | ggaaaggacgaaacaccggtCGCCCTTACATTGATCCACTTTTTTgaattctcgacctcgagaca |  |
| 28 | REX\_7 | ggaaaggacgaaacaccggtTTGACCTTGTCTCTTTTAATTTTTTgaattctcgacctcgagaca |  |
| 29 | REX\_8 | ggaaaggacgaaacaccggtCTGAGACTGTCTCTCTTTGTTTTTTTgaattctcgacctcgagaca |  |
| 30 | RIX\_1 | ggaaaggacgaaacaccggtCTGACGGTGTGCACCTCCGAATTTTTTTgaattctcgacctcgagaca |  |
| 31 | RIX\_2 | ggaaaggacgaaacaccggtGCATGACCCTACACCTACCTACTTTTTTgaattctcgacctcgagaca |  |
| 32 | RIX\_3 | ggaaaggacgaaacaccggtATGTCTCTCAATTGATTGTGTCTTTTTTgaattctcgacctcgagaca |  |
| 33 |  |  |  |
| 34 | Table 4) Ribozyme-REX/RIX-Ribozyme gblocks: | |  |
| 35 | REX1 | aactggggcacaagcttaatTAAGACTGATGAGTCCGTGAGGACGAAACGAGTAAGCTCGTCTGCCCCTTCCTTCAGACAGCGGCCGGCATGGTCCCAGCCTCCTCGCTGGCGCCGGCTGGGCAACATGCTTCGGCATGGCGAATGGGACGGCCGCaactccCACCTGCAAC |  |
| 36 | REX5 | aactggggcacaagcttaatTAAGACTGATGAGTCCGTGAGGACGAAACGAGTAAGCTCGTCCAGTCCCACCTTCCCACTTGGCCGGCATGGTCCCAGCCTCCTCGCTGGCGCCGGCTGGGCAACATGCTTCGGCATGGCGAATGGGACGGCCGCaactccCACCTGCAAC |  |
| 37 | RIX1 | aactggggcacaagcttaatTAAGACTGATGAGTCCGTGAGGACGAAACGAGTAAGCTCGTCCTGACGGTGTGCACCTCCGAATGGCCGGCATGGTCCCAGCCTCCTCGCTGGCGCCGGCTGGGCAACATGCTTCGGCATGGCGAATGGGACGGCCGCaactccCACCTGCAAC |  |
| 38 |  |  |  |
| 39 | Table 5) REX/RIX qPCR primers |  |  |
| 40 | REX\_1\_up | TGCCCCTTCCTTCAGACAGC |  |
| 41 | REX\_2\_up | GTTGAGTGAAATCAGTTTC |  |
| 42 | REX\_3\_up | TCTCACTACTCTATTATTC |  |
| 43 | REX\_4\_up | ACCCACGCCTAACTAATTC |  |
| 44 | REX\_5\_up | CAGTCCCACCTTCCCACTT |  |
| 45 | REX\_6\_up | CGCCCTTACATTGATCCAC |  |
| 46 | REX\_7\_up | TTGACCTTGTCTCTTTTAA |  |
| 47 | REX\_8\_up | CTGAGACTGTCTCTCTTTGT |  |
| 48 | RIX\_1\_up | CTGACGGTGTGCACCTCCGAAT |  |
| 49 | RIX\_2\_up | GCATGACCCTACACCTACCTAC |  |
| 50 | RIX\_3\_up | ATGTCTCTCAATTGATTGTGTC |  |
| 51 | MIR16 | TAGCAGCACGTAAATATTGGCG |  |
| 52 |  |  |  |
| 53 | Table 6 |  |  |
| 54 | exoCLIP sequencing primers: |  |  |
| 55 | Custom oligo dT-UMI primer (HPLC purified) | CAAGCAGAAGACGGCATACGAGATNNNNNNNNGTGACTGGAGTTCAGACGTGTGCTCTTCCGATCTTTTTTTTTTTTTTT |  |
| 56 | Index forward (i5) primers (TruSeq i5 primers) | AATGATACGGCGACCACCGAGATCTACAC [i5 index] ACACTCTTTCCCTACACGACGCTCTTCCGATCT |  |
| 57 | Universal reverse primer (P7) | CAAGCAGAAGACGGCATACGAG |  |
| 58 | exoCLIP sequencing barcodes: |  |  |
| 59 | MDA-U | AGGCGAAG |  |
| 60 | MDA-HNRNPA2 | TAATCTTA |  |
| 61 | MDA-HNRNPB1 | CAGGACGT |  |
| 62 | MDA-RBM24 | GTACTGAC |  |
| 63 |  |  |  |
| 64 | Table 7 |  |  |
| 65 | RT primer | GTGACTGGAGTTCAGACGTGTGCTCTTCCGATCTTTTTTTTTTTTTTT |  |
| 66 | TSO-UMI primer | iCiGCTCTTTCCCTACACGACGCTCTTCCGATCTNNNNNNrSrSrS |  |
| 67 | Takara Fwd PCR primer | AATGATACGGCGACCACCGAGATCTACAC[i5]ACACTCTTTCCCTACACGACGCTCTTCCGATCT |  |
| 68 | Takara Rev PCR primer | GATCGGAAGAGCACACGTCTGAACTCCAGTCAC[i7]ATCTCGTATGCCGTCTTCTGCTTG |  |
| 69 |  | \*\* i5 and i7 barcodes are listed in SMARTer® smRNA-Seq Kit for Illumina® User Manual cat# 635029 |  |
| 70 |  |  |  |
| 71 | Table 8) IC,CM & EV from RBP Knockdowns |  |  |
| 72 |  | Index1 | Index2 |
| 73 | IC1 | ATTCAGAA | ATAGAGGC |
| 74 | IC2 | GAATTCGT | ATAGAGGC |
| 75 | IC3 | CTGAAGCT | ATAGAGGC |
| 76 | IC4 | TAATGCGC | ATAGAGGC |
| 77 | IC5 | CGGCTATG | ATAGAGGC |
| 78 | IC6 | TCCGCGAA | CCTATCCT |
| 79 | IC7 | ATTCAGAA | CCTATCCT |
| 80 | IC8 | GAATTCGT | CCTATCCT |
| 81 | IC9 | CTGAAGCT | CCTATCCT |
| 82 | IC10 | TAATGCGC | CCTATCCT |
| 83 | IC11 | CGGCTATG | GGCTCTGA |
| 84 | IC12 | TCCGCGAA | GGCTCTGA |
| 85 | CCM1 | ATTCAGAA | GGCTCTGA |
| 86 | CCM2 | GAATTCGT | GGCTCTGA |
| 87 | CCM3 | CTGAAGCT | GGCTCTGA |
| 88 | CCM4 | TAATGCGC | AGGCGAAG |
| 89 | CCM5 | CGGCTATG | AGGCGAAG |
| 90 | CCM6 | TCCGCGAA | AGGCGAAG |
| 91 | CCM7 | ATTCAGAA | AGGCGAAG |
| 92 | CCM8 | GAATTCGT | AGGCGAAG |
| 93 | CCM9 | CTGAAGCT | TAATCTTA |
| 94 | CCM10 | TAATGCGC | TAATCTTA |
| 95 | CCM11 | CGGCTATG | TAATCTTA |
| 96 | CCM12 | TCCGCGAA | TAATCTTA |
| 97 | EV1 | ATTCAGAA | TAATCTTA |
| 98 | EV2 | GAATTCGT | CAGGACGT |
| 99 | EV3 | CTGAAGCT | CAGGACGT |
| 100 | EV4 | TAATGCGC | CAGGACGT |
| 101 | EV5 | CGGCTATG | CAGGACGT |
| 102 | EV6 | TCCGCGAA | CAGGACGT |
| 103 | EV7 | ATTCAGAA | GTACTGAC |
| 104 | EV8 | GAATTCGT | GTACTGAC |
| 105 | EV9 | CTGAAGCT | GTACTGAC |
| 106 | EV10 | TAATGCGC | GTACTGAC |
| 107 | EV11 | CGGCTATG | GTACTGAC |
| 108 | EV12 | TCCGCGAA | TATAGCCT |
| 109 | LL01A | ATCACGAT | NNNNNNNN |
| 110 | LL01B | CGATGTAT | NNNNNNNN |
| 111 | LL02A | TTAGGCAT | NNNNNNNN |
| 112 | LL02B | TGACCAAT | NNNNNNNN |
